# Sequence clustering confounds AlphaFold2

**DOI:** 10.1101/2024.01.05.574434

**Authors:** Joseph W. Schafer, Devlina Chakravarty, Ethan A. Chen, Lauren L. Porter

## Abstract

Though typically associated with a single folded state, some globular proteins remodel their secondary and/or tertiary structures in response to cellular stimuli. AlphaFold2^1^ (AF2) readily generates one dominant protein structure for these fold-switching (a.k.a. metamorphic) proteins^2^, but it often fails to predict their alternative experimentally observed structures^3,4^. Wayment-Steele, et al. steered AF2 to predict alternative structures of a few metamorphic proteins using a method they call AF-cluster^5^. However, their Paper lacks some essential controls needed to assess AF-cluster’s reliability. We find that these controls show AF-cluster to be a poor predictor of metamorphic proteins. First, closer examination of the Paper’s results reveals that random sequence sampling outperforms sequence clustering, challenging the claim that AF-cluster works by “deconvolving conflicting sets of couplings.” Further, we observe that AF-cluster mistakes some single-folding KaiB homologs for fold switchers, a critical flaw bound to mislead users. Finally, proper error analysis reveals that AF-cluster predicts many correct structures with low confidence and some experimentally unobserved conformations with confidences similar to experimentally observed ones. For these reasons, we suggest using ColabFold^6^-based random sequence sampling^7^–augmented by other predictive approaches–as a more accurate and less computationally intense alternative to AF-cluster.

## Introduction

The Paper reported successful predictions from three naturally evolved fold-switching protein families characterized previously^8^ (**Figure 1a**). Most predictions were performed on homologs of KaiB, whose fold switching helps to regulate cyanobacterial circadian rhythms. Later predictions focused on the eukaryotic mitotic spindle protein Mad2, whose N-and C-terminal regions interconvert between open and closed forms with distinct secondary structures and hydrogen bonding partners, and the transcriptional regulator RfaH, whose C-terminal domain (CTD) transforms from an autoinhibited α-helical form to an active β-sheet that binds the ribosome, fostering efficient translation.

**Figure 1.**
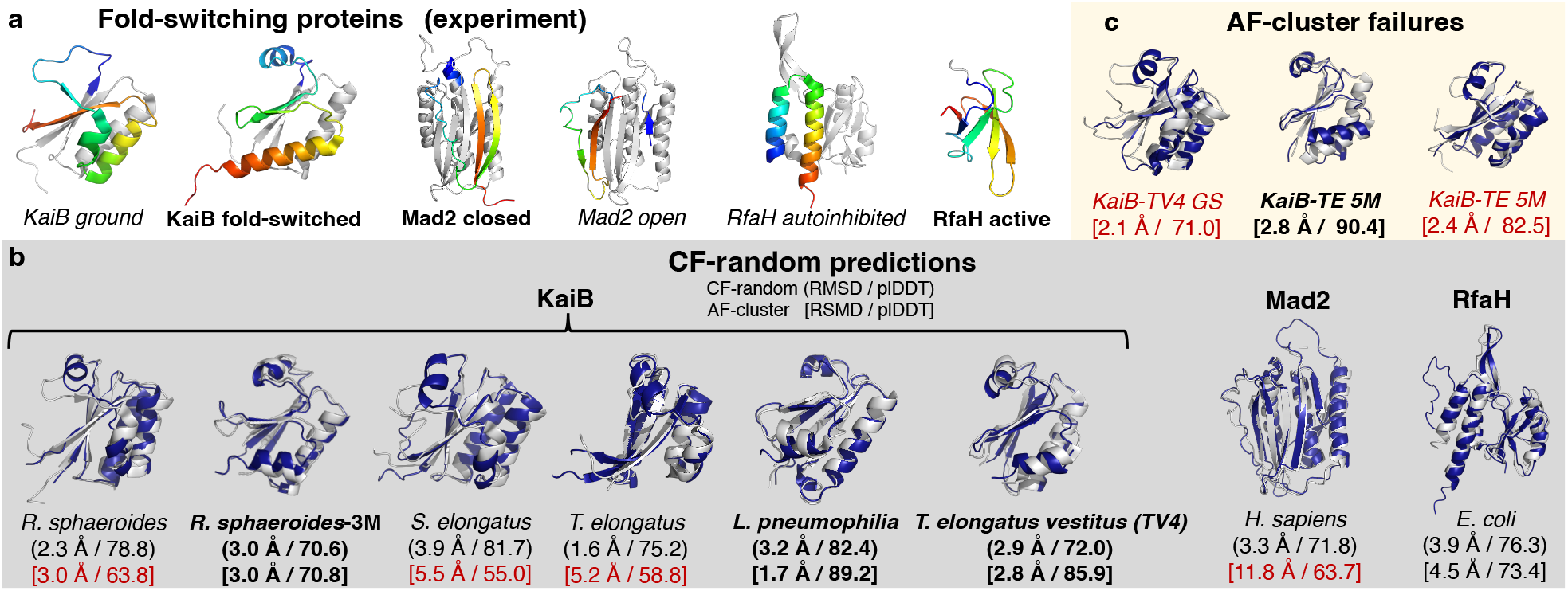
ColabFold with randomly sampled MSAs (CF-random) outperforms AF-cluster. (A) Experimentally determined structures of fold-switching proteins predicted in the Paper: KaiB, Mad2, and RfaH. Fold-switching regions are rainbow, colored N-terminus (blue) to C-terminus (red); regions that do not switch folds are gray. Full-length structures of all proteins are shown, except that the C-terminal β-sheet domain of RfaH is shown without its gray N-terminal domain to fit in the figure. Though monomeric units are shown, KaiB ground often folds into a tetramer and Mad2 forms a dimer with one of each conformer. Bold labels indicate dominant (easy-to-predict) conformations, whereas italicized labels indicate alternative. (B) Superpositions of CF-random predictions (navy) with experiment (light gray) for KaiB proteins from different organisms and for E. coli RfaH and H. sapiens Mad2. All atom root-mean-square deviations (RMSDs) and AF2 prediction confidences (plDDTs) reported below the source organisms of each protein for both CF-random (parenthesis, above) and AF-cluster (square brackets below); bold labels indicate dominant fold prediction; others are alternative. Red labels indicate RMSD > 5Å relative to experiment and/or plDDT scores <70. (C) AF-cluster mistakes two single-folding proteins as metamorphic: KaiB T. elongatus vestitus (TV4) and fold-switched stabilized KaiB from T. elongatus (KaiB-TE 5M). Though neither protein has been observed to populate the ground state (GS) fold (red font), AF-cluster predicts them with higher confidences than many of the correct predictions in (b). The correct fold-switched forms of both are predicted also; KaiB-TV4’s fold-switched form is shown in (b).

## Results

We were surprised by the Paper’s claim that sequence clusters produced higher confidence models of RfaH’s autoinhibited state than predictions from its full multiple sequence alignment (MSA). AF2’s confidence metric–the predicted local distance difference test (plDDT)–was reported as 73.9 for AF-cluster, on average, but 68.6 for full-MSA predictions. Despite our previous experience generating AF2-based predictions of RfaH^2,4,9^, we had not seen AF2 generate correct models of the autoinhibited state from full MSAs with plDDTs so low. To check, we generated 250 models by running AF2 on the full RfaH MSA from Wayment-Steele et al.’s GitHub repository (the Repo): only 2% of them had plDDT scores near the reported value (**Supplementary Figure 1a**). In fact, average predictions of RfaH’s autoinibited conformation from the full MSA (74.7) were more confident than the reported value for AF-cluster (73.9). This reported value also puzzled us. After identifying AF-cluster predictions from the Repo with helical bundle folds like the autoinhibited structure, we calculated an average plDDT of 63.8 (**Supplementary Figure 1a**). The Repo says that the plDDT score from full MSAs reported in the Paper was generated using ColabFold rather than AF2. This would seem to be an inappropriate control because the AF-cluster predictions of RfaH were generated using AF2^5^. Controls should be performed with the same software.

Since RfaH’s autoinhibited state was predicted with higher confidence from its full MSA rather than AF-cluster (**Supplementary Figure 1a**), the Paper’s claim that “clustering resulted in deconvolving conflicting sets of couplings” is called into question. Inspired by the Paper, we used MSA Transformer^10^ to assess the couplings of the sequence cluster that produced the highest confidence model of autoinhibited RfaH since it would presumably present the strongest couplings. To the contrary, no amino acid contacts unique to the autoinhibited α-helical form were observed, but weak contacts unique to the active β-sheet form were present (**Supplementary Figure 1b)**.

Further inspection of results from the Repo revealed that AF2 predictions of RfaH from uniformly sampled MSAs predicted accurate, high-confidence structures more successfully than sequence clusters (**Supplementary Figure 1c**,**d**). The authors used AF-cluster to generate 250 RfaH predictions. To assess their accuracies, we calculated the RMSDs of their fold-switching CTDs referenced against both experimentally determined conformations (**Figure 1a**): α-helical hairpin (autoinhibited) and β-sheet (active). Out of the 250 predictions, the CTDs of 10 were predicted accurately (defined as being within 3Å of either experimentally determined conformation and with an average confidence ≥ 55), a 4% success rate. By contrast, among the 21 models that AF2 generated from randomly sampled MSAs, five had accurately predicted CTDs, a 24% success rate. With both sequence clustering and random MSA sampling, AF2 predicted an incorrect hybrid α-helix/β-sheet CTD conformation (**Supplementary Figure 1c**,**d**). Nevertheless, these results suggested that better predictions may be achieved by randomly sampling MSAs with depths other than 10 or 100, the only two depths randomly sampled in the Paper.

To test whether random sequence sampling may enable robust predictions of some fold-switching proteins, we ran ColabFold^6^–an efficient-yet-accurate implementation of AF2–with randomly sampled shallow input MSAs and found that it predicted structures from all three protein families with higher accuracies and confidences than AF-cluster (**Figure 1b**). This approach, hereafter called CF-random, predicted structures from all three metamorphic protein families with higher accuracies and confidences than AF-cluster overall. Thus, randomly sampling diverse MSAs–a method proposed previously^7^–enables more accurate, higher-confidence predictions than clustering by sequence similarity. Importantly, we achieved the same outcome when we used the version of AF2 reported in the Repo: randomly sampled MSAs outperformed AF-cluster (**Supplementary Table 1**). All KaiB predictions can be reproduced from single sequences also^11^.

Though the Paper claims that AF-cluster enables predictions “scored with high confidence by AF2’s learned predicted local distance difference test (plDDT) measure,” we identified several predictions from the Repo with low plDDT scores (**Figure 1b**). Another concerning observation: AF-cluster predicts incorrect structures with relatively high confidence (**Figure 1c**). Fold-switching proteins, including KaiB and RfaH, often have single-folding homologs. Thus, a good predictor must distinguish between the two^9^. Recent work shows that AF-cluster can mispredict two single-folding RfaH homologs as metamorphic^4^. It does the same for two experimentally confirmed single-folding KaiBs (**Figure 1c**). Specifically, AF-cluster predicted that KaiB *T. elongatus vestitus* (KaiB-TV4) assumes the ground state conformation in addition to its experimentally confirmed fold-switched form. While the confidence of this ground-state prediction is higher than 4/6 correct AF-cluster KaiB predictions (**Figure 1b**), nuclear magnetic resonance (NMR) experiments from the Paper provide no evidence for it. Instead, these experiments indicate that KaiB-TV4 assumes the fold-switched conformation and a minor unfolded state^5^. By contrast, CF-random correctly predicts KaiB-TV4 to be a single folder with high confidence.

Further, AF-cluster incorrectly predicted that a fold-switch stabilized KaiB variant (KaiB-TE 5M) is metamorphic, again with higher confidence than 4/6 correct AF-cluster predictions (**Figures 1b,c**). Only five mutations distinguish this variant from its experimentally characterized ground-state homolog, but NMR experiments indicate that the variant assumes the fold-switched state only^12^. CF-random also incorrectly predicts that KaiB-TE 5M is metamorphic. These predictive errors indicate that AF2-based predictions of mutational effects should be interpreted with caution. For instance, the Paper reports a set of three mutations correctly predicted to switch the conformational balance of *R. sphaeroides* KaiB. Two of these three mutations caused AF2 to predict the same fold switch with plDDT=68.1^5^. Curiously, the Paper does not report experimental tests of this double mutant prediction, leaving us to question its accuracy.

Moving from model quality to ensemble predictions, CF-random significantly outperformed AF-cluster in both accuracy and efficiency for KaiB, Mad2, and RfaH (**Figure 2**). Ensembles predicted by CF-random and AF-cluster for *T. elongatus* KaiB and referenced against both of its experimentally determined conformations are shown in **Figure 2a**. Both methods successfully capture both conformations with high confidence. However, AF-cluster also predicts experimentally unobserved false positive conformations with high confidence, but CF-random does not. Further, CF-random predicts more true positives and fewer false positives for RfaH and Mad2 also (**Supplementary Figure 2**). Its superior accuracy is evidenced by higher Matthews Correlation Coefficients and %Success for KaiB, Mad2, and RfaH (**Figure 2b**). Furthermore, CF-random is much more efficient, requiring 1-2 ColabFold runs to generate ensembles, while AF-cluster required 95-329 AF2 runs/ensemble (**Figure 2b**).

**Figure 2.**
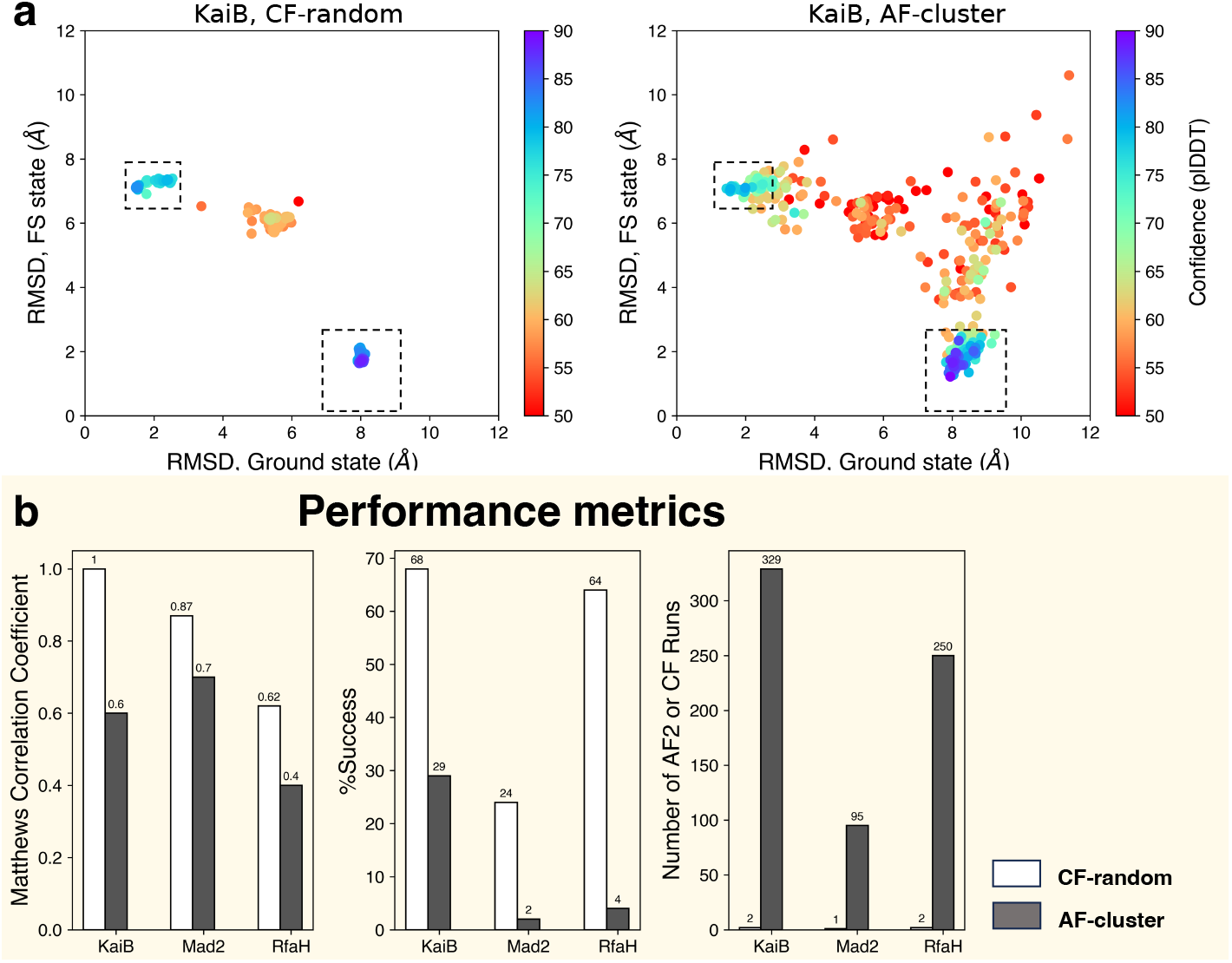
Random sequence sampling outperforms sequence clustering in the three fold-switching proteins tested in the Paper. (a). Both CF-random and AF-cluster predict both the fold-switched (lower right dashed boxes) and ground state (upper left dashed boxes) conformations of KaiB, but AF-cluster predicts high-confidence false positives (dots outside of dashed boxes with confidences ≥70). CF-random does not. Comparable numbers of predictions are compared: CF-random, 330; AF-cluster, 329. (b). CF-random has better Matthews Correlation Coefficients and %Accuracies for KaiB, RfaH, and Mad2. CF-random is also much more efficient, requiring 1-2 runs rather than 95-329 (AF-cluster).

## Discussion

These analyses indicate that CF-random, based on a method proposed previously^7^, outperforms AF-cluster, which requires more compute time and produces more false positives. However, neither CF-random nor AF-cluster predicts fold switching reliably, especially in more difficult cases. For instance, a recent benchmarking study showed that both AF2 and AF-cluster failed to predict the correct conformations of sequence-diverse RfaH homologs and two fold-switching proteins outside of AF2’s training set^4^. So does CF-random. These RfaH homologs were a more difficult test: all fold-switching sequences were <35% identical to their closest experimentally characterized homolog. For comparison, the sequences of all KaiB variants in the Paper were ≥ 47% identical to their experimentally characterized homologs. Nevertheless, the correct conformations of the more difficult RfaHs were predicted successfully using the secondary structure predictor JPred4^9,13^. As suggested by our previous work^14^, JPred4 also reproduces all of the Paper’s experimentally confirmed KaiB predictions, including their triple mutant (**Supplementary Figure 3**). Thus, JPred4 could be used to augment AF2-based predictions of difficult fold switchers that undergo α-helix <-> β-sheet transitions. Further, coevolutionary signatures corresponding to both conformations of fold-switching proteins can be used to confirm correctly predicted alternative folds^15^. In short, AF2 is an outstanding tool for generating three-dimensional models of protein structure, but its current ability to accurately predict alternative conformations is limited. We suggest pairing its predictions with orthogonal validation–such as JPred4 predictions and/or coevolutionary signatures–to enhance predictive accuracy.

## Methods

CF-random was run with ColabFold1.5.3 with depths max-seq = 1, 8, 64 for KaiB, Mad2, and RfaH, respectively, and max-extra-seq = 2*max-seq in all 3 cases. All other parameters were kept constant. More details about predictions and other calculations can be found in **Supplementary Methods**.

## Data

Code and results can be found at https://github.com/porterll/CF-random.

## Acknowledgements

L.L.P thanks Gabriel Rocklin, Loren Looger, Marius Clore, Eugene Koonin, Gisela Storz, Susan Marqusee, George Rose, Brian Volkman, Aaron Robinson, Jaime Fraser, Nick Fawzi, and Myeongsang Lee for helpful discussions. This work utilized resources from the NIH HPS Biowulf cluster (http://hpc.nih.gov), and it was supported by the Intramural Research Program of the National Library of Medicine, National Institutes of Health (LM202011, L.L.P.).

## Notes

https://github.com/porterll/CF-random

### Competing Interest Statement

The authors have declared no competing interest.

### Summary of Updates

Supplement with more methodological details.

