## Supplemental Methods and figures for "Sequence clustering confounds AlphaFold2"

Supplementary information for:
Sequence clustering confounds AlphaFold2

Joseph W. Schafer[1], Devlina Chakravarty[1], Ethan A. Chen[1], and Lauren L. Porter[1,2,*]

[1]National Center for Biotechnology Information, National Library of Medicine, National Institutes of Health, Bethesda, MD 20894
[2]Biochemistry and Biophysics Center, National Heart, Lung, and Blood Institute, National Institutes of Health, Bethesda, MD, 20892

### Supplementary Methods

*RfaH structure comparisons*
AlphaFold2.2.0[1] was used to generate 250 structures of *E. coli* RfaH using RfaH's ColabFold MSA from the Repo. All 5 AF2 models were calculated with the deterministic flag turned on; all other AF2 settings were default including 3 recycles. AF-cluster models were taken directly from the Repo, and the root mean square deviations (RMSDs) of their CTDs (residues 115-147) were compared with an experimentally determined structure of autoinhibited RfaH (PDB ID: 2OUG) using Cealign in PyMOL[16]. The full structures of all models with RMSDs ≤ 5Å were then realigned to 2OUG using align in PyMOL with no outlier rejection. Models with RMSDs ≤ 5Å were curated manually and can be found on the CF-random Github repo. Average plDDT scores of these remaining structures were calculated directly from their PDB files; plDDTs of the AF2.2.0 structures were extracted from their ranking_debug.json files.

*RfaH couplings*
With default settings, MSA Transformer[10] was used to calculate the residue-residue couplings of the best performing sequence cluster for the autoinhibited form of RfaH, RfaH_049.a3m. This was determined to be the best performer because it had the highest plDDT score among all experimentally consistent autoinhibited predictions.

*Structure predictions*
CF-random was run using ColabFold1.5.3[6] with 16 seeds, 5 structures/seed, and max-seq = 1, 8, 64 for KaiB, Mad2, and RfaH, respectively, and max-extra-seq = 2*max-seq in all 3 cases. All structures in Figure 1b were generated using these methods. To determine whether the superior performance of CF-random was due to ColabFold, we ran AlphaFold2.2.0 on all KaiB sequences, *E. coli* RfaH, and *H. sapiens* Mad2 with the same max-seq and max-extra-seq specifications as CF-random, 3 recycles, 5 models/protein. This approach, AF-random, performed similarly to CF-random (**Table S1**). Structures of KaiB-TE 5M were generated using the 2QKEE_colabfold.a3m MSA from the Repo, since the sequences of 2QKE and KaiB-TE 5M are 95% identical. We generated clusters and structure predictions using the AF-cluster Colab notebook. As suggested by the site, model 3 was used. Model 1 and 2 predictions crashed. Since a full MSA for KaiB-TV4 was not provided on the Repo (none was needed for their calculations), we generated one using Colabfold and inputted it into the AF-cluster Colab notebook to get sequence clusters. Since the Repo states that KaiB-TV4 was predicted using Colabfold with 12 recycles, we ran ColabFold on all clusters with 12 recycles, Model 3.

*Experimental comparison*
RMSDs in **Figure 1** were calculated using PyMOL align with no outlier rejection. Mad2, RfaH, and KaiB proteins were referenced against their corresponding experimentally determined structures with two exceptions. Since no structures of its ground-state form have been deposited in the PDB, KaiBRS was referenced against the its closest experimentally determined homolog assuming the ground state conformation: 4KSO, and KaiBRS_3m was referenced against 8FWY, chain M – KaiBRS in the fold-switched form.

*Ensemble generation*

The CF-random *T. elongatus* KaiB ensemble was generated by running ColabFold1.5.3 with 33 seeds, 5 structures/seed in two separate runs: one with max-seq = 1, max-extra-seq =2, the other without max-seq and max-extra-seq specifications. Similarly, the CF-random *E. coli* RfaH ensemble was generated by running ColabFold1.5.3 with 25 seeds, 5 structures/seed in two separate runs: one with max-seq = 32, max-extra-seq =64, the other without max-seq and max-extra-seq specifications. The CF-random *H. sapiens* Mad2 ensemble was generated by running ColabFold1.5.3 with 19 seeds, 5 structures/seed in one run: max-seq=8, max-extra-seq=16. RMSDs in **Figure 2 and Supplementary Figure 2** were calculated with MDTraj[17] on alpha carbons only. Full RMSDs were used for KaiB and Mad2 and average plDDTs were required to be ≥70 for accuracy calculations. RMSDs of RfaH's C-terminal domain (CTD, resis 115-162) were used because of its flexible linker, which prevents accurate whole-protein comparisons. Average plDDTs of RfaH's CTD were required to be ≥55 since predictions of the α-helical CTD conformation are less certain than the NTD and β-sheet CTD conformation.

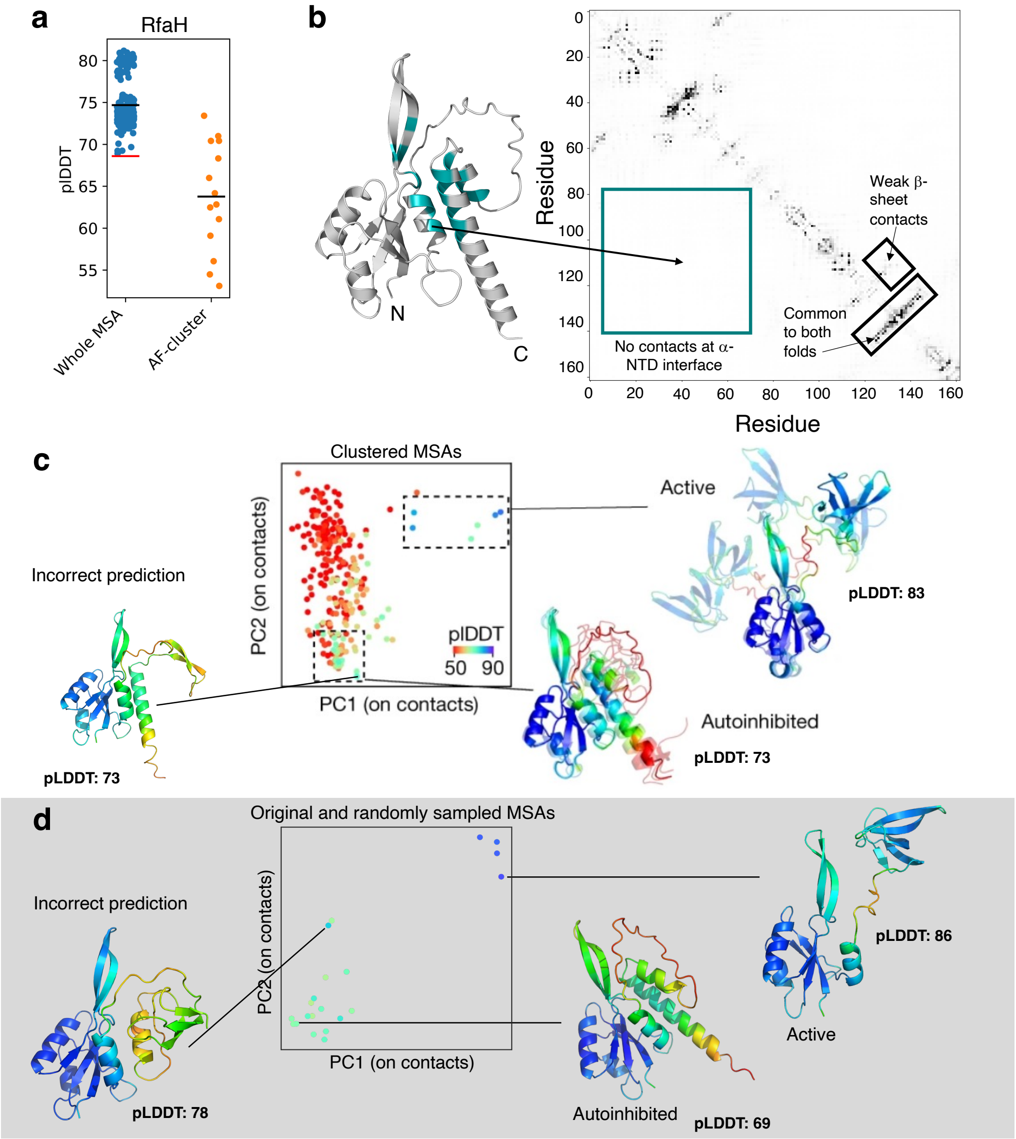

*Figure 1. For RfaH, accurate than sequence clustering. (a). AF2 predicts the autoinhibited conformation of RfaH with more confidence from a whole MSA than from random sequence sampling. Black lines represent mean values for whole MSAs (74.7) and AF-cluster (63.8). Red line is the plDDT value that the Paper uses to represent plDDT values from whole MSAs (68.6). (b) MSA Transformer predicts no unique coevolutionary signals from shallow MSAs used to predict the autoinhibited structure of RfaH. Contacts unique to this state is highlighted in teal. Black boxes in the RfaH contact map are not represented in the structure and are used instead to annotate interesting features of its contact map. (c) AF-cluster predicts both conformations of RfaH with reasonably high confidence. It also predicts an incorrect hybrid α/β conformation with equivalent confidence. Overall accuracy: 10/250 predictions assumed either the autoinhibited or active conformations with prediction confidences of fold-switching C-terminal domains ≥ 55. Plot and left protein structures taken directly from the Paper (Figure 4b). (d) Random sequence sampling also produces both conformations with higher accuracy using the same confidence thresholds: 5/21 models assume experimentally consistent conformations. Hybrid α/β structures are also predicted, though with lower confidence in the fold-switching C-terminal domain compared to AF-cluster. Plots generated using code and data from the Repo. RfaH structures also from the Repo.*

**Table 1**

| Protein | Method | Ranked[1,2] | RMSD (Å)[1] | plDDT |
|---|---|---|---|---|
| KaiB-Rs | AF-cluster | WP_069333443 | 3 | 63.8 |
| KaiB-Rs | CF-random | 1 | 2.3 | 78.8 |
| KaiB-Rs | AF-random | 1 | 2.2 | 71.6 |
| KaiB-Rs-3M | AF-cluster | Single sequence | 3 | 70.8 |
| KaiB-Rs-3M | CF-random | 1 | 3 | 70.6 |
| KaiB-Rs-3M | AF-random | 1 | 2.9 | 70 |
| KaiB-Se | AF-cluster | WP_011242647 | 5.5 | 55.0 |
| KaiB-Se | CF-random | 1 | 3.9 | 81.7 |
| KaiB-Se | AF-random | 0 | 4.7 | 78.5 |
| KaiB-Te | AF-cluster | WP_011056333 | 5.2 | 58.8 |
| KaiB-Te | CF-random | 1 | 2.3 | 75.2 |
| KaiB-Te | AF-random | 1 | 2.3 | 72.7 |
| KaiB-Lp | AF-cluster | WP_027265801 | 1.7 | 88.9 |
| KaiB-Lp | CF-random | 1 | 3.2 | 82.4 |
| KaiB-Lp | AF-random | 2 | 2.7 | 82.6 |
| KaiB-TV4 | AF-cluster | WP_024125951 | 2.8 | 85.9 |
| KaiB-TV4 | CF-random | 1 | 2.9 | 72.0 |
| KaiB-TV4 | AF-random | 2 | 3.7 | 56.7 |
| Mad2 | AF-cluster | 1S2H_047 | 11.8 | 63.7 |
| Mad2 | CF-random | 20 | 3.3 | 71.8 |
| Mad2 | AF-random | 4 | 10.4 | 65.7 |
| RfaH | AF-cluster | RfaH_049 | 4.5 | 73.4 |
| RfaH | CF-random | 34 | 3.9 | 76.3 |
| RfaH | AF-random | 1 | 3.8 | 74.9 |

[1]AF2 Rankings of CF- and AF-random predictions are specified. Since AF-cluster produces 1 structure/MSA (Model 1), the MSA used to generate the prediction is reported.
[2]ColabFold's best ranked structure is 1; AlphaFold's is 0. This ranking is maintained in the table. Note that all CF-random predictions use the structure ranked 1, except for Mad2, whose CF-random predictions produce higher ranked structures in the open state. The top ranked structure in the closed state is 20.
[3]Root mean square deviations (RMSDs) for each prediction (rankings left) are reported. All RMSDs were calculated using PyMOL align with no outlier rejection, except for KaiB-Rs, whose closest experimentally determined homolog with the same fold was 4KSO. In this case, PyMOL's Cealign was used instead since it gives more accurate RMSDs for less similar sequences.

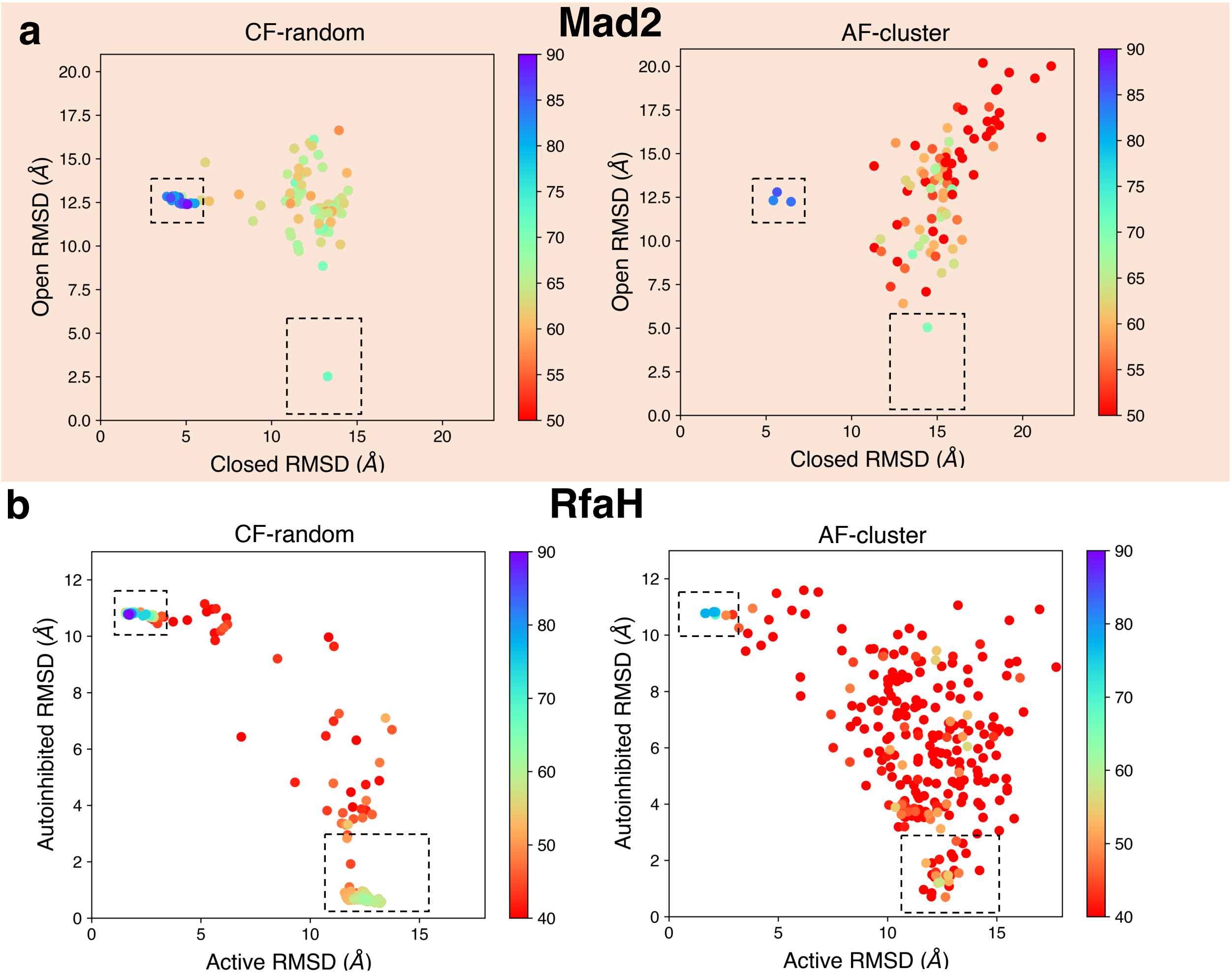

*Figure 2. As with KaiB, CF-random predicts both conformations of Mad2 and RfaH with higher accuracy than AF-cluster. (a) CF-random predicts both conformations of Mad2 (dotted boxes) with a 24% success rate while AF-cluster predicts them with 2% success. Predictions in boxes were counted as successful if the average plDDT score of the whole protein exceeded 70. Both plots show 95 predictions total. (b). CF-random predicts both conformations of RfaH (dotted boxes) with a 64% success rate while AF-cluster predicts them with 4% success. Predictions in boxes were counted as successful if the average plDDT score of the C-terminal domain (residues 115-162) exceeded 55. Both plots show 250 predictions total.*

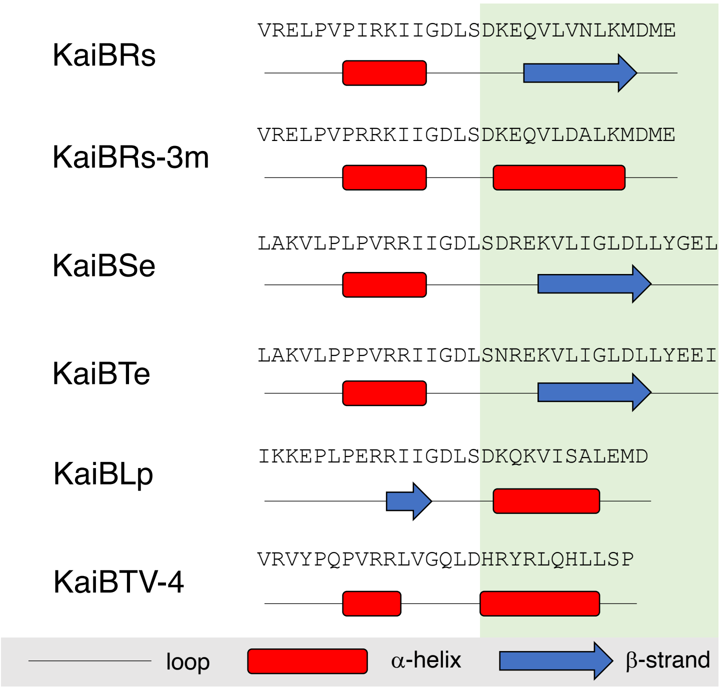

*Figure 3. Secondary structure predictions can be used to accurately infer fold switching in experimentally characterized KaiB proteins. Wayment-Steele et al. use the C-terminal secondary structure element from AF2 predictions to infer whether a KaiB variant assumes the ground (β-strand) or fold-switched (α-helix) state. The same can be done here: the C-terminal secondary structure elements (green background) correspond correctly to experimentally determined ground (β-strand) and fold-switched (α-helix) states. The same predictions were used to accurately infer fold-switching in the RfaH family of transcriptional regulators in 10/10 cases. AF-cluster predictions were successful in 4/10. Thus, secondary structure predictions sometimes offer equal or more predictive insights than AF2.*
